# Neurons in the ventral striatopallidal complex modulate lateral hypothalamic orexin/hypocretin neuron activity: Implications for reward-seeking

**DOI:** 10.1101/2020.03.05.979468

**Authors:** Caitlin S. Mitchell, Aida Mohammadkhani, Elizabeth E. Manning, Erin J. Campbell, Simon D Fisher, Jiann W. Yeoh, Amy J. Pearl, Nicholas J. Burton, Min Qiao, Jacqueline A. Iredale, Jaideep S. Bains, Gavan P. McNally, Zane A. Andrews, Brett A. Graham, Thomas E. Scammell, Bradford B. Lowell, Dong Kong, Stephanie L. Borgland, Christopher V. Dayas

## Abstract

Reward-seeking involves the engagement and computation of multiple physiological and motivational parameters. The lateral hypothalamus (LH) is a necessary node in the circuits that control food-seeking and motivation. One group of cells that plays an important yet incompletely understood role in these processes are the orexin/hypocretin (OX/HT) neurons. OX/HT cells are located exclusively within the LH and are implicated in feeding, arousal, and reward-seeking behavior. Importantly, the role of OX/HT neurons in consummatory versus reward-seeking actions is not fully defined, nor are the circuits that control the activity of these neurons under different behavioral states. Here we show that OX/HT neurons respond in real time to food presentation and that this response is modulated by differences in metabolic state. We observed increased OX/HT neuron activity on approach to food, but this activity trended towards pre-approach levels by the start of the consummatory phase. Next, we studied ventrostriatopallidal (VSP) inputs to the OX/HT neurons. Using optogenetics and cell type-specific electrophysiology, we found that ventral pallidum inputs onto OX/HT neurons exert strong inhibitory (and weak excitatory) effects whereas the lateral nucleus accumbens shell provides weaker direct inhibitory connections with OX/HT neurons. These findings reveal that the activity of OX/HT neurons is strongly modulated by metabolic and hedonic state. Further, OX/HT neurons is primarily associated with food approach and that the effect of VSP-terminal output is to suppress OX/HT activity.

## Introduction

Orexin/hypocretin (OX/HT) neurons are located exclusively within the lateral hypothalamus (LH) with widespread projections to cortical and subcortical structures (De Lecea et al., 1998) (Peyron et al., 1998) (Sakurai et al., 1998). Early work identified a role for OX/HT neurons in feeding and sleep/wake behavior (Espana et al., 2002) (Sakurai et al., 1998). Indeed, seminal studies firmly implicated OX/HT in food consumption with intracerebroventricular injections of OX/HT increasing food intake and body weight (Sakurai et al., 1998). Systemic treatment with an OX/HT-1 receptor antagonist prevented feeding behavior (Haynes et al., 2000). Despite this, more recent studies have demonstrated that the role of OX/HT in consummatory behavior is not so straightforward and may be influenced by changes in metabolic state. For example, ablation of OX/HT neurons using genetically targeted diphtheria toxin increased body weight in mice (Ramanathan & Siegel, 2014). Additionally, measures of *in vivo* cell activity also demonstrated that OX/HT cells are rapidly inactivated by eating, irrespective of caloric content or metabolic state (González et al., 2016).

Whilst there is a clear, historical link between the OX/HT system and consumption, recent work supports a broadened role for these neurons in general arousal-related behaviors, including avoidance to aversive stimuli (Viviani et al., 2015). For example, OX/HT cell activity is strongly increased by negatively valanced emotional stimuli such as predator odor (Giardino et al., 2018). There is also a large literature linking the OX/HT system with drug motivated behavior, including stress- and cue-evoked drug-seeking as well as opiate withdrawal-related behaviors (Borgland et al., 2009) (Boutrel et al., 2005) (Campbell et al., 2020) (Harris et al., 2005) (Georgescu et al., 2003) (James et al., 2011). Thus, previous literature has linked OX/HT cells with both consumption and the motivation for reward. Still, new neuroscience technology affords us a more nuanced examination of the involvement of OX/HT neurons in food approach versus consumption. This information is vital to understand the pathogenesis of reward-related disorders such as obesity and substance use.

To resolve the continued uncertainty surrounding the role of the OX/HT system in approach versus consummatory actions, we assessed the reactivity of OX/HT neurons to palatable or normal food using *in vivo* fiber photometry under different metabolic states. These recordings revealed three key findings. First, that GCaMP6f-derived activity signals of OX/HT neurons were associated with reward approach and began to decline, trending towards baseline during consumption. Second, under food restriction the presentation of normal chow induced a significantly larger activity response than in *ad libitum* fed mice, as did normal chow following injection of ghrelin. Third, the presentation of a palatable chocolate reward to fed mice increased OX/HT activity to levels identical to food restricted mice.

We then considered what brain circuits might be responsible for modulating OX/HT cell activity. Traditional anatomical work and more recent retrograde monosynaptic viral tracing studies have shown that the ventrostriatopallidal complex (VSP), including the nucleus accumbens (NAc) and ventral pallidum (VP), sends significant anatomical projections to intra-lateral hypothalamic (LH) circuits with the potential to control OX/HT activity (Giardino et al., 2018) (Zhang & Kelley, 2000). Regarding consummatory and approach behaviors, previous work has shown that activation of NAc shell (NAcSh) terminals in the LH suppresses alcohol seeking behavior and food consumption (Gibson et al., 2018) (O’Connor et al., 2015). However, O’Connor et al. (2015) found no evidence of synaptic connectivity with OX/HT neurons. Further, Sheng and colleagues found that NAc core projections to the LH disinhibit LH-glutamate neurons (including OX/HT) through a polysynaptic pathway involving a local GABAergic relay (Sheng et al., 2023). Based on these mixed findings, we re-investigated how VSP inputs to the LH influence OX/HT neurons using optogenetics and cell type-specific electrophysiology to understand the specific contributions of the VSP and NAc to these effects. Together, our data indicate that the activity of OX/HT neurons is primarily associated with food approach and that VSP output decreases OX/HT activity.

## Results

### Metabolic state and palatability can enhance the activity of OX/HT cells

To examine the temporal profile of OX/HT cells and how metabolic state may influence this activity, orexin-Cre mice were injected with Cre-dependent AAV5-GCaMP6f virus into the perifornical-lateral hypothalamic area (herein referred to as LH) and implanted with a fiber optic cannula 0.2 mm dorsal to the LH (**Fig. 1A**). In mice used to verify the specificity of the virus, GCaMP6f labelling was visualized in the LH consistent with the known distribution of OX/HT neurons (Peyron et al., 1998). Importantly, 97% (±0.9%) of all GCaMP6f transduced cells were dual labelled for orexin-A and 53.5% (±3.6%) of OX/HT-positive LH cells were GCaMP6 positive (**Fig. 1B-C**), demonstrating that our mouse line can selectively express GCaMP6f in OX/HT neurons.

**Figure 1.**
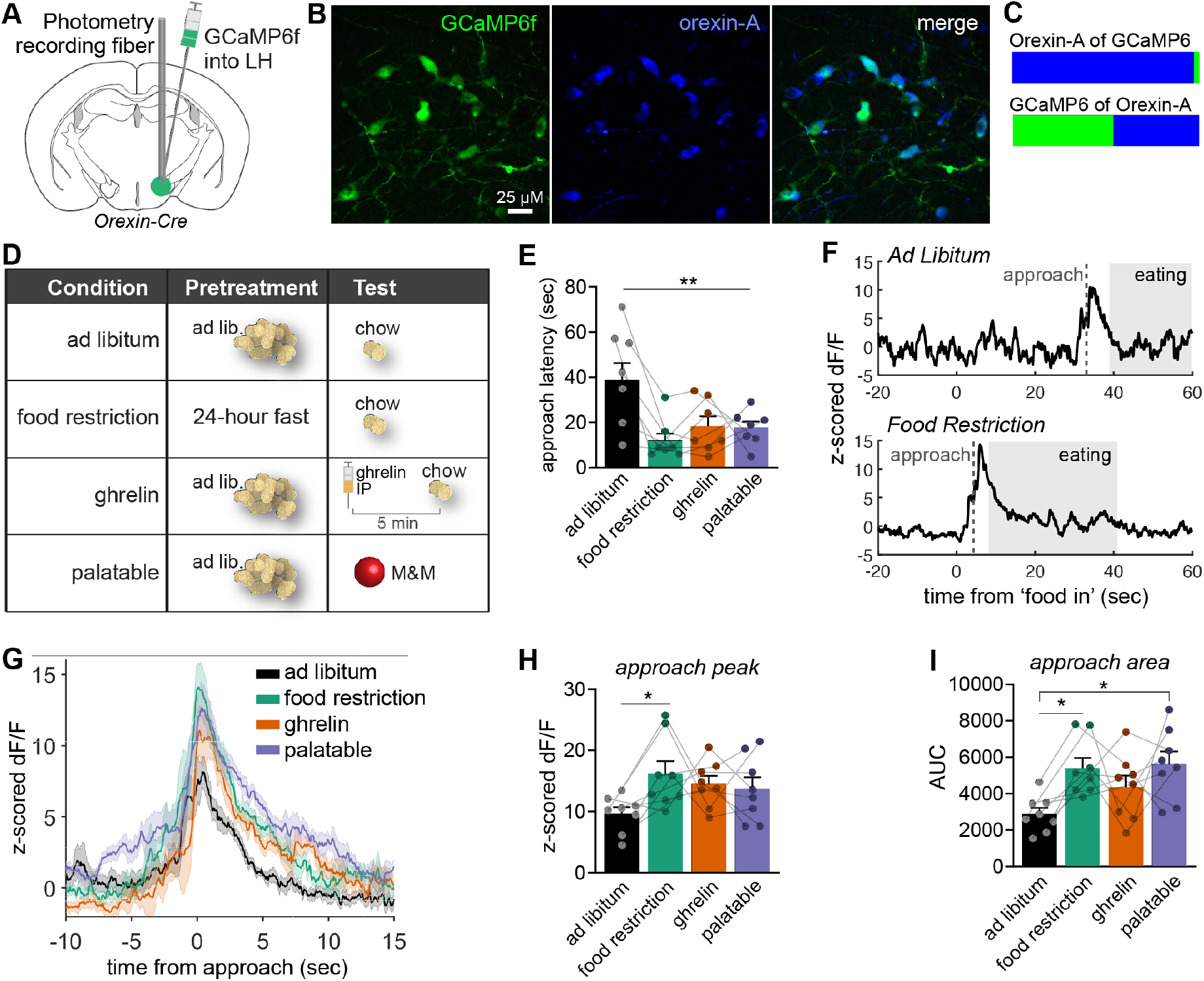
Lateral Hypothalamic (LH) orexin (OX/HT) neurons respond on approach to food objects, modulated by metabolic state. **(A)** Orexin-Cre mice received LH-directed injections of DIO-GCaMP6f and implantation of a fiber optic probe. **(B)** Microscopy images showing orexin-Cre dependent GCaMP6f expression in the LH, together with immunolabelling of orexin-A positive cells, and a merged image. **(C)** Proportion of GCaMP6+ cells that are Orexin-A+ (97% ±0.9%, n = 4 mice), and Orexin-A+ cells that are GCaMP6+ (53.5% ±3.6%). **(D)** Experimental groups, with their pretreatments and test configurations. **(E)** The latency to approach the food pellet was significantly different between groups. Post-hoc comparisons revealed that each group had a significantly lower approach latency compared to *Ad Libitum* (p’s < 0.05). **(F)** Example z-scored dF/F photometry traces for the *Ad Libitum* and Food Restriction conditions. **(G)** Mean z-dF/F traces and shaded standard error of the mean (SEM) region for each condition, centered around the time of first approach to food. **(H)** The peak amplitude of the z-dF/F signal during the approach period varied significantly by treatment condition with post-hoc comparisons showing a significant difference between *Ad Libitum* and Food Restriction (p = 0.048). **(I)** Area under the curve (AUC) analysis of the approach period also significantly varied by treatment condition, with post-hoc comparisons showing significant differences between *Ad Libitum* and Food Restriction (p = 0.039), and *Ad Libitum* and Palatable conditions (p = 0.021). Error bars = +SEM; scale bars: 25μm; ^*^p<0.05; ^**^p<0.01.

To study Ca^2+^ based activity dynamics in OX/HT neurons under different metabolic states, orexin-Cre::DIO-GCaMP6f mice were tested with the presentation of standard food (chow) placed into their home cage under three metabolic conditions (**Fig. 1D**) – *ad libitum* fed + normal chow (*Ad Libitum* condition); 24 h fasted + normal chow (Food Restriction condition); *ad libitum* fed + injections of the appetite promoting hormone + normal chow (Ghrelin condition). We also investigated the approach to a palatable chocolate food pellet in *ad libitum* fed mice (a mini chocolate M&M™, Palatable condition). Mice were exposed to all conditions in a counterbalanced design. A significant reduction in latency to approach the food was found in all conditions, in comparison to the *Ad Libitum* fed condition (F_3,18_ = 7.419, p = 0.002; Bonferroni post-hoc comparisons p’s<0.05) (**Fig. 1E**). Latency to approach in the Food restricted, Ghrelin injected, and Palatable conditions did not differ (p’s>0.05). During approach to the test food item, Ca^2+^ activity in OX/HT neurons increased from baseline in all groups, irrespective of metabolic state (examples **Fig. 1F**; mean signals **Fig. 1G**). The peak amplitude of the z-dF/F signal during the approach period (from first approach to start eating) varied significantly by condition (F_3,21_ = 3.103, p = 0.049), with post-hoc comparisons revealing a significant increase in Food Restriction peak amplitude compared to *Ad Libitum* (p = 0.048) (**Fig. 1H**). Additionally, there was a significant difference across treatment conditions in area under the curve (AUC) of the Ca^2+^ signal during approach (F_3,21_ = 4.510, p = 0.014). Post-hoc comparisons revealed significant increases in AUC of the Ca^2+^ signal during approach in the Food restriction group (p = 0.039) and Palatable group (p = 0.021) versus *Ad Libitum* controls (**Fig. 1I**).

Together these data suggest both metabolic state and palatability can enhance the activation of OX/HT cells. Importantly, a close inspection of the temporal profile of OX/HT cell activity indicated that GCaMP6f fluorescence was closely linked to food approach rather than consummatory actions – the signal peaked during approach, prior to food consumption, and trended towards baseline levels during consumption (**Fig. 1F**).

### The ventrostriatopallidal complex makes monosynaptic inhibitory projections to OX/HT cells

To understand what brain circuits might be responsible for modulating OX/HT activity, we first verified that TdTomato fluorescence driven by AAV8-*h-orexin-tdTomato* was limited to LH OX/HT neurons (**Fig. 2B**). To test if VP neurons are functionally connected with LH OX/HT neurons, we injected ChR2 into the VP of VGAT-Cre mice and recorded optically evoked currents from TdTomato fluorescent LH OX/HT neurons (**Fig. 2A-B**). Of 13 recorded neurons, we evoked monosynaptic currents in 12 neurons identified by the remaining current in the presence of tetrodotoxin (TTX) and 4-aminopyridine (4-AP). The averaged optically evoked IPSC amplitude was 153 ± 24pA (**Fig. 2C-D**) with an average latency of 11 ± 1.5ms (**Fig. 2E**), suggesting that almost all inhibitory projections from the VP to LH OX/HT neurons are monosynaptic. Because some VP neurons are also thought to co-express excitatory vesicular transporters (Root et al., 2018; Tooley et al., 2018) we tested if VP GABAergic neurons could also release glutamate (**Fig. 2F**). We found that optical stimulation of VP GABAergic terminals in the LH could produce either monosynaptic (23 ± 8pA, 5 neurons) or polysynaptic excitatory postsynaptic currents (EPSCs) (35 ± 12pA, 2 neurons) with a latency of 13 ± 2ms or 20 ± 0.5ms, respectively (**Fig. 2G-H**). Importantly, the amplitude of oEPSCs was substantially smaller than oIPSCs. Taken together, these data suggest that under specific recording conditions, VP GABAergic neurons can release both GABA and glutamate onto LH OX/HT neurons, and inputs to the LH from VP GABA neurons are primarily inhibitory.

**Figure 2.**
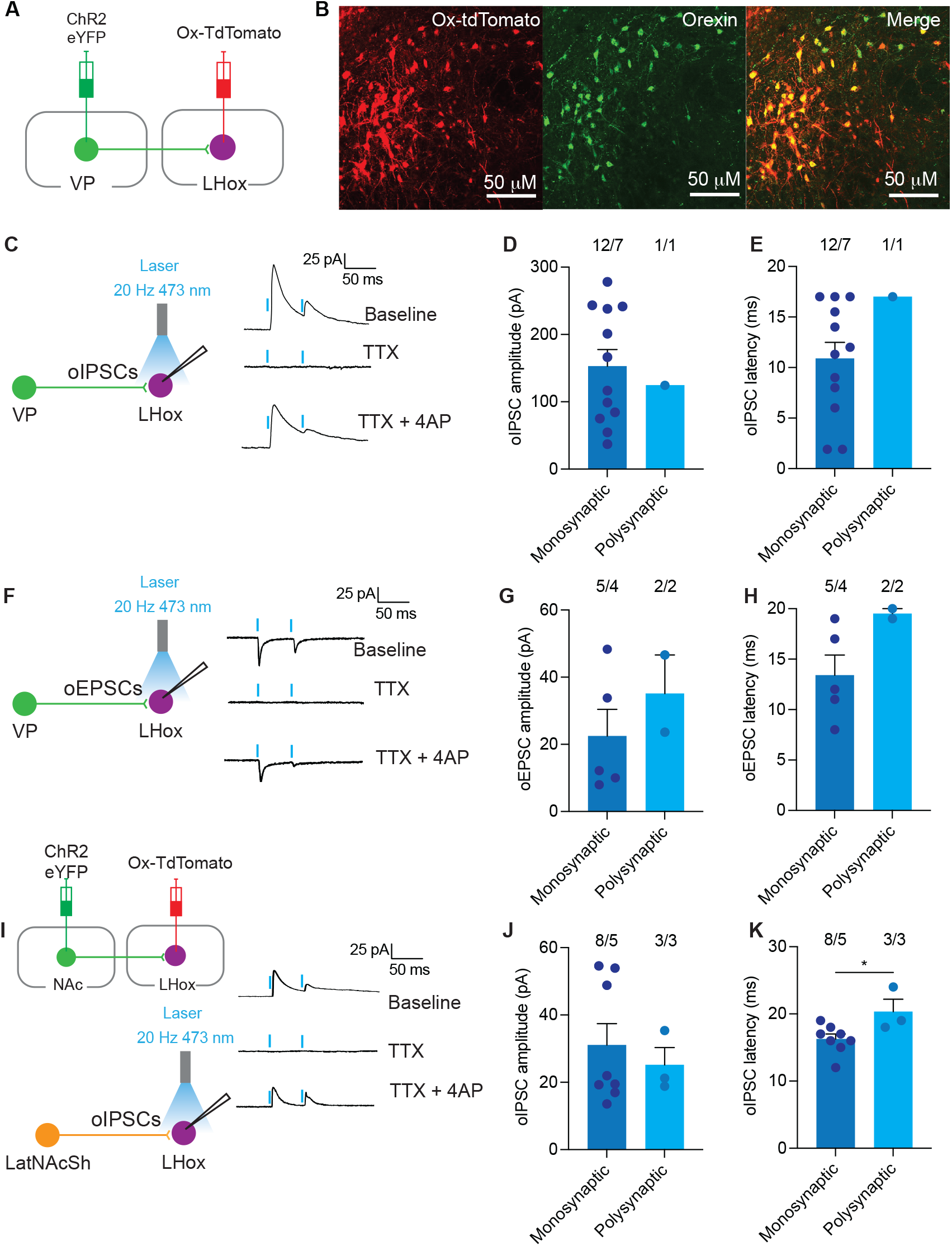
The ventral pallidum and the nucleus accumbens lateral shell make monosynaptic inhibitory projections to lateral hypothalamic (LH) orexin (OX/HT) neurons. **(A)** Cre-dependent ChR2 was expressed in the ventral pallidum (VP) of VGAT-Cre mice and TdTomato was expressed in LH OX/HT neurons. **(B)** Example image of TdTomato expressing OX/HT neurons (red), orexin neurons labelled with Alexa 488 (green), and the merged image. **(C)** Example traces of optically evoked inhibitory postsynaptic currents (IPSCs) onto LH OX/HT neurons during baseline, during TTX (1μM) application and then during TTX + 4-AP (100μM) application. **(D)** A comparison of monosynaptic and polysynaptic current amplitude (n/N = 12 cells/7 mice). Only one polysynaptic current was detected suggesting that VP projections to LH OX/HT neurons are primarily monosynaptic. **(E)** Latency from optical stimulation to oIPSC peak. **(F)** We tested if it was possible to evoke optical EPSCs from VP GABAergic terminals onto LH OX/HT neurons. Example traces of optically evoked EPSCs onto LH OX/HT neurons during the baseline, TTX, and TTX + 4-AP application. **(G)** A comparison of monosynaptic (n/N = 5 cells/4 mice) and polysynaptic current (n/N = 2 cells/2 mice) amplitude. **(H)** Latency from optical stimulation to oEPSC peak. **(I)** Cre-dependent ChR2 was expressed in the lateral shell of the nucleus accumbens of VGAT-Cre mice and TdTomato was expressed in LH OX/HT neurons. Example traces of optically evoked IPSCs onto LH OX/HT neurons during baseline, TTX, and TTX + 4-AP application. **(J)** A comparison of monosynaptic (n/N = 8 cells/5 mice) and polysynaptic current (n/N = 3 cells/3 mice) amplitude. **(K)** Latency from optical stimulation to oIPSC peak. The latency of polysynaptic currents was significantly longer compared to monosynaptic currents (p = 0.033). Error bars = +SEM

We next tested if the NAcSh sent monosynaptic projections to LH OX/HT neurons by injecting a Cre-dependent ChR2 to the medial or lateral shell of the NAc of VGAT-Cre mice (**Fig. 2I**). We found that medial shell injections did not produce any optically evoked IPSCs or EPSCs. However, lateral shell NAc projections produced either monosynaptic (31 ± 6pA, 8 neurons) or polysynaptic (25 ± 5pA, 3 neurons) optically evoked IPSCs with latencies of 16 ± 0.8ms or 20 ± 2ms, respectively (t(9) = 2.5, p = 0.033) (**Fig. 2J-K**). We were unable to optically evoke EPSCs in the medial shell to LH OX/HT projection. Taken together, these data suggest that lateral but not medial shell NAc neurons make inhibitory projections to LH OX/HT neurons.

## Discussion

We used a combination of circuit dissection techniques including channel rhodopsin-assisted circuit mapping and cell type-specific fiber photometry to establish how LH OX/HT cells are modified by changes in metabolic state. Our data indicates that OX/HT activity peaked on approach to food but declined on arrival at the food source and trended towards baseline during consummatory behavior. Channel rhodopsin-assisted circuit mapping revealed that LH OX/HT neurons receive substantial monosynaptic inhibitory input from the VP. Together these data suggest that VP inputs likely inhibit LH OX/HT cell activity and may gate or limit reward-seeking.

### Orexin (OX/HT) cell activity is associated with approach to food

Using fiber photometry targeted at LH OX/HT cells, we showed that this population of neurons primarily responds to food approach. GCaMP6f fluorescence declined following approach to food, which is consistent with previous research that implicated LH OX/HT neurons in reward-seeking and motivation for sweet food (Harris et al., 2005) (Thorpe et al., 2005), and increased arousal and locomotor behavior (Alexandre et al., 2013). It has also been shown that food restricted mice exhibit increased c-Fos activity in OX/HT neurons following food anticipation (Mieda et al., 2004). Notably, we found limited evidence for changes in Ca^2+^ fluorescence during food consummatory actions. This finding appears to contrast earlier work linking LH OX/HT with increases in food intake (Sakurai et al., 1998). However, more recent work has suggested that the increases in food consumption may be secondary to enhanced behavioral arousal and locomotion, such as that produced by high LH OX/HT peptide injections (Blais et al., 2017) (Sutcliffe & de Lecea, 2002) (Nevarez & de Lecea, 2018) (Zink et al., 2018). In keeping with this line of thinking, the deletion of LH OX/HT cells using viral-mediated diphtheria toxin increased food consumption and weight gain (González et al., 2016).

A report studying the Ca^2+^ activity of LH OX/HT neurons found that activity rapidly declined after contact with foods of different tastes, textures, and nutrient content (González et al., 2016). Indeed, the authors argued that the act of eating rapidly inactivates OX/HT neurons. This interpretation was supported by the timescale of effects of food contact or licking on Ca^2+^ activity dynamics in OX/HT neurons making direct inhibitory actions of nutrients (such as glucose) on OX/HT cell firing unlikely. In the present study we did not detect a rapid reduction in activity in OX/HT cells during eating, regardless of metabolic state or palatability, but rather the reduction in OX/HT cell activity was more closely correlated with the termination of approach behavior.

### Lateral hypothalamic orexin (LH OX/HT) activity during approach to food is modulated by metabolic state and salience

Interestingly, LH OX/HT neurons displayed greater Ca^2+^ activity depending on metabolic state. For example, food restriction significantly increased Ca^2+^ activity to standard chow. Enhanced OX/HT signaling in these conditions is consistent with work of (Horvath & Gao, 2005) who showed that food restriction was sufficient to increase the frequency of miniature EPSCs and increase Vglut2-positive puncta onto OX/HT neurons. One interpretation of these findings is that food restriction increases OX/HT excitatory drive to increase food-seeking and approach behaviour. Notably, we also found that injections of ghrelin tended to elevate OX/HT cell activity to levels seen under food-deprivation, even though the mice were allowed free access to food. Ghrelin is a peptide secreted by the stomach that is thought to stimulate feeding behavior through actions on growth hormone secretagogue receptor (GHSR)-expressing neurons in the arcuate nucleus (Hewson et al., 2002). Ghrelin can also increase OX/HT activity in electrophysiological slice or dispersed cell preparations, indicating possible direct effects in the LH (Kohno et al., 2003). We also found that the LH OX/HT cell responses to palatable food reached levels equivalent to when mice had been food restricted. Thus, not only does metabolic state influence LH OX/HT cell activity but so does approach to a more salient food object.

Our work is also consistent with behavioral studies showing that orexin receptor antagonists reduce motivated responding for palatable food (Borgland et al., 2009) and drugs of abuse under high effort progressive ratio conditions (James et al., 2011). Furthermore, drug-induced increases in excitatory drive to LH OX/HT neurons promotes reward-seeking behavior but has little impact on drug consumption (Bentzley & Aston-Jones, 2015). Together, this data suggests that the LH OX/HT system acts to enhance effort-based approach behavior according to the valance of the stimulus and that the level of activity can be updated through actions from metabolic signals and synaptic drive.

### The ventral striatopallidal (VSP) complex provides inhibitory inputs to orexin (OX/HT) neurons

The termination of activity after food approach raised a key question as to the possible brain regions that might inhibit OX/HT activity. We focused on VSP, given prior work showing that inhibitory inputs from the ventral striatum can terminate feeding-related behaviors and alcohol-seeking (Gibson et al., 2018) (O’Connor et al., 2015) and the finding that the VSP provides strong direct anatomical projections with OX/HT neurons (Giardino et al., 2018). Using Vgat-Cre mice and virally mediated expression of the reporter protein tdTomato, we found evidence of mono and polysynaptic inhibitory connections to LH OX/HT cells. These findings provide the first evidence of functionally patent connections from the VSP to these neurons. Importantly, while we also observed some evidence of oEPSCs, currents evoked under these conditions were substantially weaker.

The present study confirms and extends recent work demonstrating that medial NAcSh D1R spiny projection neurons make little to no contact onto OX/HT (or melanin concentrating hormone)-expressing cells in the LH (O’Connor et al., 2015). Consistent with this, our findings revealed a lack of connectivity between these regions. Interestingly however, lateral shell injections did reveal connectivity to LH OX/HT neurons, in this case but both mono and polysynaptic inhibitory currents were observed.

The precise connectivity within the LH that permits indirect polysynaptic inhibitory actions remains to be fully understood but is likely a consequence of the complex local LH circuitry that gates the input to OX/HT neurons. For example, OX/HT neurons receive input from local excitatory and inhibitory interneurons (Ferrari et al., 2018) (Xie et al., 2006). Further work will be required to determine whether the indirect inhibition of OX/HT cells involves disinhibition of a local GABAergic microcircuit or inhibition of local glutamatergic interneurons.

### Conclusions

Overall, the present study demonstrates that the activity of LH OX/HT neurons precedes food consumption and is tightly associated with appetitive approach behavior rather than consummatory actions. The activity of LH OX/HT neurons is also enhanced by metabolic state and as well as the salience of food. Therefore, OX/HT neurons appear involved in appetitive approach behavior, independent of actual food intake. Electrophysiological patch-clamping of LH OX/HT neurons demonstrated that the VSP provides inhibitory signals to LH OX/HT cells. Future studies will need to determine exactly how these pathways control LH microcircuits to gate approach and other behaviors. Given the work of studies such as Giardino and colleagues (2018) showing that OX/HT neurons are also robustly activated by aversive stimuli, we hypothesize that disruption of food approach, such as through engagement with a predator might usurp this inhibitory pathway. Understanding how these pathways are disrupted under pathological states such as obesity or substance use disorder may highlight new insights into treating these conditions using behavioral or pharmacological interventions.

## Methods

### Lead contact and material availability

This study did not generate new unique reagents. Further information and requests for resources and reagents should be directed to and will be fulfilled by the Lead Contact, Chris Dayas.

### Ethics statement

For photometry experiments, all procedures were performed in accordance with the Prevention of Cruelty to Animals Act (2004), under the guidelines of the National Health and Medical Research Council (NHMRC) Australian Code of Practice for the Care and Use of Animals for Experimental Purposes (2013). Animal ethics were approved by The University of Newcastle Animal Ethics Committee. For electrophysiology experiments, all protocols were in accordance with the ethical guidelines established by the Canadian Council for Animal Care and were approved by the University of Calgary Animal Care Committee.

### Mice

For photometry experiments, male and female knock-in orexin-IRES-Cre (orexin-Cre) mice (n=4 for viral validation; n=8 for photometry) were obtained from BIDMC Neurology (Harvard Medical School) (De Luca et al., 2022) (Xiao et al., 2021). Mice were hemizygous and single housed. For electrophysiology experiments, 21 male and female Vgat-ires-cre mice (VGATcre; Slc32a1tm2(cre)Lowl/J) were obtained from Jackson Laboratory (strain: 016962) and bred in the Clara Christie Centre for Mouse Genomics. Mice were group housed in groups of 3-5. Food and water were available *ad libitum* and all mice were maintained on a reverse 12-hour light/dark cycle (0700 lights off).

The following conditions were used in photometry experiments:

*Ad libitum*: mice were allowed *ad libitum* access to chow leading up to testing. The test food was a normal chow pellet.

*Food restriction*: mice were not allowed access to their home caged chow for 24 hours prior to testing. The test food was a normal chow pellet.

*Ghrelin IP*: mice were allowed *ad libitum* access to chow leading up to testing. The test food was a chow pellet.

*Palatable food*: mice were allowed *ad libitum* access to chow leading up to testing. The test food was a mini chocolate M&M.

### Surgery

To validate the selectivity of viral expression in orexin-Cre mice and to conduct fibre photometry experiments, mice were anesthetized with isoflurane (5% induction, 2% maintenance) before being placed into a stereotaxic frame (Stoelting Instruments). Each mouse received unilateral Cre-dependent viral injections of AAV5-CAG-FLEX-GCaMP6f (Addgene) directed at the LH (0.3μl; anteroposterior (AP): -1.4, mediolateral (ML): ±1.0, dorsoventral (DV): -5.25). The virus was injected at a rate of 0.15μl/min using a 2μl Hamilton Neuros syringe (30G) attached to a Stoelting Quintessential Stereotaxic Injector pump. For photometry experiments, fiber optic probes (6mm; 400μm core; Metal Ferrule Cannula (MF2.5); 0.37NA; Doric Lenses, QC, CAN) were inserted just above the LH (AP: -1.4, ML: ±1.0, DV: -5.1).

For electrophysiology experiments, vgat-Cre mice were anesthetized with isoflurane (vapourized at 3-5%, with O_2_ at 0.5-1L/min), injected with subcutaneous meloxicam (5mg/kg) and saline (2-3mL, 0.9%), and secured in the stereotaxic frame (Kopf; Tujunga, CA). Channelrhodopsin (ChR2) (AAV2-EF1a-DIO-hChR2(H134)-EYFP; Neurophotonics, Centre de Recherche CERVO, Quebec City, QC, Canada) was bilaterally infused into either the ventral pallidum (VP; AP: -0.15, ML: ±1.5, DV: -5) nucleus accumbens lateral shell (latNacSh; AP: -1.42, ML: ±1.75, DV: -4.90) or nucleus accumbens medial shell (medNacSh; AP: +1.42, ML: ±0.85, DV: -4.90) at a volume of 300nl/hemisphere and a rate of 60nl/min with an additional 10 min before raising the nanoinjector. Mice were returned to home cages for 4 to 5 weeks for full virus expression before use in experiments. Two weeks prior to electrophysiology experiments, mice were injected with AAV8-hSyn-orexin-tdTomato into the LH (0.2μl; AP: -1.35, ML: ±0.9, DV: -5.25) to visualize OX/HT neurons in the LH. This virus was diluted 1:4 in sterile saline solution before stereotaxic injection.

### Fiber Photometry

To visualize Ca^2+^ fluorescence using single fiber photometry; a PC running Doric Lenses Software (Doric Neuroscience Studio version 5.3.3.1) was connected to a Fiber Photometry Console (FPC; Doric Lenses, QC, CAN). The FPC was attached via a BNC to a 4-channel LED Driver. A 465nm wavelength LED (CLED; Doric Lenses) was used to stimulate GCaMP and reflected light was detected in the 525nm spectrum using a filter cube consisting of dichroic mirrors (6-port Fluorescent Mini Cube with an integrated photodetector head; iFMC; Doric Lenses). A 405nm LED (CLED; Doric Lenses) was used as an isosbestic control, as fluorescence detected from this wavelength is not Ca^2+^ dependent, so hence changes in signal can be attributed to autofluorescence, bleaching and fiber bending. Light from these fiber-coupled LEDs entered the filter cube via FCM-FCM patch cords (200μm core; 0.22NA; Doric Lenses). Light exited the filter cube via a FC-MF2.5 patch cord (400μm core; 0.48NA; Doric Lenses). Mice were attached to this patch cord via the implanted fiber optic cannula with cannula and patch cord communicating via a plastic mating sleeve (Thorlabs). Emission light from the mouse is transmitted back up through the same patch cord. A lock-in amplifier (Doric Lenses) was used to modulate the signals. Power output from the end of the FC-MF2.5 patch cord was determined using a photodiode power meter (Thorlabs, NJ, USA). Power output from both LEDs was set at approximately 40uW, with power never exceeding 60uW.

Mice were habituated in their home cage to the test set-up for 3 days prior to testing (15mins/day; including patch cord connection). Each mouse underwent the four separate test conditions: Ad libitum, Food restriction, Ghrelin IP, and Palatable. The tests were counterbalanced for mice to account for order effects. Test days were separated from each other by 2 or 3 days. For Ad libitum, Food restriction and Palatable food groups, mice were first attached to the patch cord and habituated for 5 minutes and GCaMP6f fluorescence recorded for 5 minutes. At the 5-minute mark, the appropriate food pellet was placed at the opposite side of the home cage to where the mouse was located. For the Ghrelin IP group, mice were given a 5-minute baseline recording and a saline control injection to account for any hyperarousal effects of IP injection. 10 minutes later, mice were injected with 0.3mg/kg of ghrelin and 5 minutes later a normal chow pellet was placed in the home cage.

### Photometry Analysis

Doric photometry traces were synchronized with the live video recording (Blackfly S GigE, FLIR systems, USA) in order to determine event times: approach to food, begin eating, stop eating. The demodulated output from two channels (isosbestic control 405nm and Ca^2+^-dependent 465nm wavelengths) was exported from the Doric Neuroscience Software. All analysis was performed using custom MATLAB scripts. 465nm and 405nm signals were first averaged with a sliding window, then baseline correction was performed using the adaptive iteratively reweighted penalized least squares method (Zhang et al., 2010). Next, robust z-scores were calculated for both signals ((signal – median) / median absolute deviation). The isosbestic signal was then aligned to the Ca^2+^-dependent signal using linear regression (MATLAB’s lasso function). Finally, the fitted isosbestic signal was subtracted from the Ca^2+^-dependent signal to give the z-score dF/F values. The peak z-dF/F value during the approach was measured as the maximum value during the period from 5 seconds prior to ‘first approach’ identified by video observation, until the ‘begin eating’ event. This maximum value was baseline corrected with a baseline measure comprising the mean of 5 seconds of activity prior to food presentation. Area under the curve was measured using the MATLAB trapz function, of the same time period.

### Slice Electrophysiology recordings

Mice were deeply anaesthetized with isoflurane and transcardially perfused with an ice-cold N-methyl-D-glucamine (NMDG) solution containing (in mM): 93 NMDG, 2.5 KCl, 1.2 NaH2PO4.2H2O, 30 NaHCO3, 20 Hepes, 25 D-glucose, 5 sodium ascorbate, 3 sodium pyruvate, 2 thiourea, 10 MgSO4.7H2O, 0.5 CaCl2.2H2O and saturated with 95% O2-5% CO2. Mice were then decapitated, and the brain extracted. Horizontal midbrain sections (250 μm) containing the LH were cut on a vibrating-blade microtome (Leica, Nussloch, Germany). Slices were then incubated in NMDG solution (32°C) and saturated with 95% O2-5% CO2 for 10 min. Following this, the slices were transferred to artificial cerebrospinal fluid (aCSF) containing (in mM): 126 NaCl, 1.6 KCl, 1.1 NaH2PO4, 1.4 MgCl2, 2.4 CaCl2, 26 NaHCO3, 11 glucose, (32°C) and saturated with 95% O2-5% CO2. Slices were incubated for a minimum of 45 min before being transferred to a recording chamber and superfused (2 mL min-1) with aCSF (32-34DC) and saturated with 95% O2-5% CO2. Neurons were visualized on an upright microscope using ‘Dodt-type’ gradient contrast infrared optics and whole-cell recordings were made using a MultiClamp 700B amplifier (Molecular Devices, San Jose, CA, USA). The recorded signal was collected at a sampling rate of 20 kHz and a 2 kHz Bessel filter was applied to the data during collection. OX/HT neurons in the LH were identified by red fluorescence. Recorded neurons were positioned so that the soma was at the centre of the objective. For optically evoked currents, a 5 ms, 3.5 mW, pulse of 20 Hz 470 nm was provided by an LED light source through a NA 40/0.80 water immersible Olympus microscope objective. For voltage clamp recordings, recording electrodes (3-5 MOhms) were filled with cesium methanesulfonate (CsMeSO4-Cl) internal solution containing (in mM): 85 CsCH3SO3, 60 CsCl, 10 HEPES, 0.2 EGTA, 1 MgCl2 5 QX-314-Cl, 2 MgATP, 0.3 NaGTP. Optically evoked currents were recorded at +10 mV for IPSCs and -65 mV for EPSCs in the presence of TTX (1 μM) followed by TTX and 4-AP (100 μM) to determine monosynaptic connection between the VP GABA neurons and LH OX/HT neurons. Electrophysiological data were acquired and analysed using Axon pClamp v11.2.

### Immunohistochemistry

Mice were deeply anesthetized with isoflurane and transcardially perfused with phosphate buffered saline (PBS) and then with 4% paraformaldehyde (PFA). Brains were dissected and post-fixed in 4% PFA at 4°C overnight, then switched to 30% sucrose. Frozen sections were cut at 30μm using a cryostat. For validation of orexin-Cre mice, 3 LH sections per mouse were selected and incubated in an orexin-A rabbit polyclonal antibody (1:1000 anti-rabbit, Phoenix Pharmaceuticals, CA, USA) for 48 hours at room temperature. Brain sections were then washed in 0.1M PBS to rinse unbound antibody and moved to a secondary antibody (1:500, Alexa Fluor AMCA anti-rabbit; Jackson Immunoresearch, PA, USA) for 2 hours at room temperature. Tissue slices were then mounted and cover-slipped using a mixture of 0.1M PBS and glycerol. Images of tissue sections were made using Olympus CellSens Software (version 1.3) on a microscope (Olympus BX51) at 10x objective using the green and blue wavelength filters. Virally transduced GCaMP6 cells and orexin-A labelled cells (Bregma -1.22 to -1.7) were counted using the Cell Counter add-on application in ImageJ, and analysis was undertaken by an observer blind to the treatment. To examine colocalization of TdTomato and orexin in VGATcre mice, 10% goat serum was applied to block non-specific binding for 1 hour. Coronal sections were then incubated with primary antibody rabbit orexin 1:500 (Phoenix pharmaceuticals, H-003-30) and chicken red fluorescent protein (RFP) 1:2000 (Rockland, 600-901-379) to amplify the orexin and TdTomato signal in 1% BSA for 24 hours at 40°C followed by incubation with secondary antibody Alexa Fluor 488 goat anti rabbit 1:400, Alexa Fluor 594 goat anti-chicken 1:400, and DAPI 1:2000 for 1 hour. All images were obtained on a NIKON A1R multiphoton microscope with a 20x objective (Nikon Canada, Ontario, Canada). Images were processed using ImageJ/FIJI.

### Statistical analysis

Statistical analyses were conducted using JASP V16.2 and GraphPad Prism 10. Repeated measures one-way ANOVAs with bonferroni post-hoc comparisons were performed to compare experimental conditions in the photometry experiments. For latency to approach photometry analyses, one mouse was excluded as it had a latency to approach that was more than two standard deviations above the mean. For electrophysiology, an independent samples t-test was used to examine the difference in IPSC latencies between monosynaptic and polysynaptic currents in lateral shell NAc projections to LH OX/HT neurons. Statistical significance was determined by a threshold of p<0.05.

## Funding and Disclosure

All authors declare no conflict of interest.

This work was supported by grants to CVD from the NHMRC 2016/GNT1125478; 2018/GNT1165679 and ARC DP220102567 Australia.

## Notes

### Competing Interest Statement

The authors have declared no competing interest.

### Summary of Updates

We have updated our author list, as previous contributors were inadvertently left off. We have also performed additional electrophysiological experiments with our collaborator Dr Stephanie Borgland's team to resolve the contributions of the ventral striatopallidal complex to hypothalamic orexin/hypocretin neuron activity. These findings resolved that the ventral palladium provides strong inhibitory (and weaker) excitatory input to orexin/hypocretin neurons that within the accumbens, inhibitory inputs arise from the lateral shell.

